# SILICA: Streamline Independent Component Analysis for Trajectory-Resolved White Matter Decomposition

**DOI:** 10.64898/2026.08.06.743368

**Authors:** Lei Wu, Vince D. Calhoun

## Abstract

Whole-brain tractography reconstructs the major white matter pathways as millions of individual streamlines, offering an exceptionally rich description of neural geometry. Yet the statistical methods used to compare these reconstructions across individuals inevitably discard key information. Voxel-based analyses sacrifice pathway continuity, trajectory-based methods rarely support population-level statistical decomposition, and connectome models largely abstract away the underlying geometry. No existing framework jointly characterizes the population-level statistical organization of white matter and the three-dimensional geometry of the pathways from which that organization is expressed.

We introduce streamline independent component analysis (SILICA), a framework that links group-level voxel-space statistical decomposition to subject-specific trajectories through a sparse streamline-by-voxel fingerprint. Each streamline is represented by its physical path length within a common anatomical voxel grid while retaining an explicit index-level link to its original trajectory. A two-stage dimensionality reduction reconciles tractograms of differing size and enables continuous component loadings to be back-reconstructed for every original streamline. These subject-specific loadings support weighted trajectory visualization and can be projected into voxel space to generate track-weighted component maps for conventional image-based visualization and future voxel-wise analysis. Separately, the learned group spatial components can be expressed on an independently reconstructed representative whole-brain tractogram to generate a compact trajectory-resolved atlas for group-level visualization. SILICA is a single decomposition expressed simultaneously in statistical and geometric form.

SILICA was evaluated in diffusion MRI tractograms from 30 healthy adults. The recovered spatial patterns correspond to recognizable commissural, projection, and association systems. Back-reconstructions preserved individual trajectory variation while isolating components shared across the group, and their projection into voxel and trajectory space yielded interpretable maps and atlases. As a proof of concept, SILICA has not yet been validated against anatomical reference standards or evaluated for reproducibility and performance relative to established methods. Nevertheless, these results establish a coherent foundation for analyzing white matter in a framework that jointly represents population-level statistical structure and streamline geometry.

## 1. Introduction

Diffusion-weighted magnetic resonance imaging (dMRI) and tractography have transformed the study of structural brain connectivity in vivo by providing unprecedented access to the organization of white matter pathways (Basser, Pajevic, Pierpaoli, Duda, & Aldroubi, 2000; Behrens et al., 2003; Jeurissen, Descoteaux, Mori, & Leemans, 2019). Tractography reconstructs putative fiber trajectories from local diffusion information and allows investigators to examine the structural pathways that support cognition, behavior, and neurological function (Catani & Thiebaut de Schotten, 2008; Johansen-Berg & Behrens, 2009). A typical whole-brain tractogram, however, contains hundreds of thousands to millions of streamlines without anatomical labels or an intrinsic organizational structure. Transforming this collection into an interpretable representation of white matter anatomy remains a central challenge in diffusion neuroimaging.

The objective is not merely to assign each streamline to a predefined bundle. A useful tract representation should preserve several complementary forms of information: where a tract is located, how its streamlines traverse the brain, how strongly individual streamlines contribute to it, and how these properties vary across subjects. Such features are relevant to the study of individual differences, disease-related alterations, neurosurgical planning, and the development of population-level white matter atlases. A representation that retains only bundle labels, regional connection strengths, or voxelwise density cannot fully describe the geometry and subject-specific organization of the underlying trajectories.

Existing approaches have broadly followed two main strategies, each with notable limitations. Streamline-based fiber clustering methods group individual trajectories according to geometric similarity and have produced anatomically meaningful white matter parcellations (Garyfallidis et al., 2018; L. J. O’Donnell & Westin, 2007; Zhang et al., 2018). These methods retain the trajectories assigned to each cluster, but establishing consistent group-level correspondence requires additional registration, atlas construction, or bundle-recognition procedures, because streamline geometry and sampling vary across subjects (Garyfallidis et al., 2018; Zhang et al., 2018). Many conventional approaches also assign streamlines to discrete clusters, which may not fully represent peripheral, transitional, or overlapping trajectories (L. J. O’Donnell & Westin, 2007; Zhang et al., 2018). Scalar measures such as fractional anisotropy (FA), streamline length, and other geometric or microstructural features are typically quantified only after bundles have been identified and therefore do not contribute directly to the tract-parcellation objective. (Colby et al., 2012; Yeatman, Dougherty, Myall, Wandell, & Feldman, 2012; Yeh, 2020). Consequently, the resulting group-level representation is determined primarily by streamline geometry and does not directly model graded streamline contributions, scalar information, and subject-specific variation within a unified decomposition.

Connectome-based approaches provide a different representation by summarizing tractography as a matrix in which nodes correspond to predefined regions and edges represent estimated connection strengths (Hagmann et al., 2008; Sporns, Tononi, & Kötter, 2005). This abstraction supports graph-based analysis but reduces the contributing streamlines to region-to-region connections (Hagmann et al., 2008; Lauren J O’Donnell, Golby, & Westin, 2013). Two sets of streamlines may connect the same pair of regions while following different courses through the white matter. Once those trajectories are represented by a single edge value, their curvature, spatial extent and internal course are no longer explicitly represented and cannot be recovered from the connectivity matrix alone (Lauren J O’Donnell et al., 2013).

Methods based on predefined tract definitions provide another route for subject-level bundle identification and quantitative analysis. Automated fiber quantification (AFQ) identifies established white matter fascicles from whole-brain tractography using anatomical constraints and samples diffusion or microstructural measures along each bundle to generate tract profiles (Colby et al., 2012; Yeatman et al., 2012). Supervised approaches such as TractSeg instead predict a predefined set of tract masks directly from fiber-orientation information and can subsequently support bundle-specific tractography and tractometry (Wasserthal, Neher, & Maier-Hein, 2018). These methods provide efficient and anatomically standardized representations of known pathways, but their bundle definitions are specified through anatomical rules, atlases, or training labels. They therefore serve a different purpose from data-driven decomposition, in which tract components and their streamline contributions are estimated directly from whole-brain data without a fixed tract atlas.

Track-density imaging (TDI) and the broader family of track-weighted imaging (TWI) methods provide complementary voxelwise representations of tractography-derived information. These methods project streamline-derived quantities into image space and can represent streamline density, path length, orientation, or associated scalar measures at high spatial resolution (Calamante, 2017; Calamante, Tournier, Smith, & Connelly, 2012). They provide detailed spatial summaries but do not themselves decompose whole-brain tractograms into group-level components or establish component-wise correspondence with the contributing subject-specific streamlines.

Our previous work introduced connectivity matrix independent component analysis (cmICA) as a data-driven alternative (Wu & Calhoun, 2023; Wu, Calhoun, Jung, & Caprihan, 2015; Wu, Duda, Iraji, & Calhoun, 2025). cmICA constructs a whole-brain voxel-by-voxel structural connectivity matrix in which each entry represents tractography-derived connectivity between two voxels. Group independent component analysis decomposes this matrix into spatial source maps and corresponding connectivity profiles. The resulting components localize primarily within white matter and correspond to recognizable tract systems, demonstrating that ICA can recover meaningful white matter organization from tractography-derived connectivity without predefined regions or tract labels. While the decomposition itself is informed by the geometry, the voxel-by-voxel representation of each component, however, does not retain the original streamline paths. Once the tractogram is converted into a connectivity matrix, the spatial course of each contributing streamline cannot be reconstructed, and subject-specific differences in curvature, bundle extent, branching, and trajectory dispersion become inaccessible. The matrix can also grow extremely large, increasing storage and computational demands. These limitations motivated a representation that preserves the group-level spatial decomposition achieved by cmICA while maintaining a direct link to the original streamlines.

Here we introduce SILICA, or streamline ICA, a framework that decomposes whole-brain tractograms into group-level spatial components while retaining each streamline’s original trajectory. SILICA operates directly on a streamline-level representation of each tractogram, from which group ICA yields spatial components in voxel space together with continuous component loadings for the original streamlines. Since these loadings are estimated at the level of individual streamlines, each group-level spatial component remains explicitly linked to the subject-specific streamlines that contribute to it. Unlike hard streamline clustering, SILICA permits graded streamline contributions rather than requiring exclusive assignment to a single bundle. In contrast to voxel- or connectivity-matrix-based decomposition, it retains a direct link between each spatial component and the original subject-specific streamlines. Because the original streamline coordinates are retained, component-specific tractograms can be recovered directly for each subject, and the spatial sources remain defined in a common voxel space that supports group-level estimation and anatomical interpretation across subjects. The name SILICA evokes silica, the material used to guide optical signals through fibers, reflecting the framework’s purpose of transforming an unstructured tractogram into coherent components.

In this proof-of-concept study, we first introduce the SILICA framework by comparing it with the conventional cmICA and describing its mathematical formulation, streamline representation, individual- and group-level workflows, and back-reconstruction strategy. We then apply SILICA to MNI-aligned whole-brain tractograms from 30 healthy adults, yielding group-level spatial tract source maps in voxel space and continuous loadings that quantify each streamline’s contribution to each component. Following this, we examine the resulting tract patterns at the group level and map the component loadings back to the original tractograms to construct subject-specific, streamline-resolved tract bundles. Using 12 subjects as illustrative examples, we demonstrate that shared group components can be linked to subject-specific trajectories while preserving visible intersubject differences in tract geometry, extent, density, and dispersion. We further show how the recovered streamline loadings can be projected into track-weighted component maps for downstream statistical voxel analysis. Separately, the learned group spatial source maps are mapped to generate a compact trajectory atlas. Together, these analyses show that SILICA can provide a soft decomposition of whole-brain tractograms into anatomically coherent group-level components while linking them directly to subject-specific bundles that preserve streamline details. This study establishes the feasibility of trajectory-resolved group ICA and provides a foundation for the broader validation and benchmarking that remain as future work.

## 2. Methods

### 2.1 SILICA Framework

Conventional cmICA represents a tractogram through voxel-to-voxel connectivity, such that both the observations and the resulting component profiles are defined in connectivity voxel space. This representation summarizes connections between voxel pairs but does not preserve the identity of the individual streamlines that contribute to each connection. SILICA instead represents each tractogram as a sparse streamline-by-voxel track-fingerprint matrix *M_s_*, in which each row corresponds to one original streamline and each column corresponds to one voxel in a common template space. The resulting representation preserves the correspondence between each matrix row and its original subject-specific trajectory. Importantly, ICA is performed on voxel-space fingerprints rather than directly on raw streamline coordinates. Trajectory resolution is retained because each fingerprint row remains indexed to one original streamline throughout dimensionality reduction and back-reconstruction.

SILICA decomposes whole-brain tractograms into group-level spatial components while retaining an explicit correspondence between those components and the original subject-specific streamlines. For K components, SILICA models the track-fingerprint matrix of subject *S* as

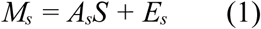

where *M_s_* ∈ ℝ*^Ns^*^×*V*^, *S* ∈ ℝ*^K^*^×*V*^is the shared spatial component matrix, *A_s_* ∈ ℝ*^Ns^*^×*K*^ is the subject-specific streamline loading matrix, and *E_s_* denotes residual variation. Here, *N_s_* is the number of streamlines for subject *s*, and *V*is the number of voxels in the common template grid. Each row of *S* defines a spatial component over the common voxel grid, whereas the corresponding column of *A_s_* quantifies the continuous expression of that spatial component by every original streamline from subject *s*. Specifically, *a_sik_* denotes the loading of streamline *i* on component *k* from subject *s*.

This formulation permits graded streamline contribution rather than requiring an exclusive bundle assignment and preserves an explicit link between each group-level spatial component and its contributing subject-specific trajectories. A streamline may also contribute to multiple components with different loading values. Because the rows of *M_s_* retain their correspondence to the original tractogram, the estimated loadings can be mapped directly back to the original subject-specific trajectories without geometric averaging or fixed-point streamline resampling.

Figure 1 summarizes the relationship between cmICA and SILICA and the extension of SILICA to group estimation. Panel A illustrates the voxel-by-voxel connectivity representation used by cmICA. Panel B shows the streamline-by-voxel representation used by SILICA and the resulting decomposition into spatial source maps and per-streamline loadings. Panel C presents the group-level workflow. Tractograms are first transformed into a common template space, and each subject-level track-fingerprint matrix is reduced independently while retaining the voxel dimension. The reduced subject matrices are then combined, reduced at the group level, and decomposed using spatial ICA to estimate shared spatial components. The group- and subject-level reduction operators are subsequently propagated backward to recover subject-specific component loadings at the original streamline resolution.

**Figure 1.**
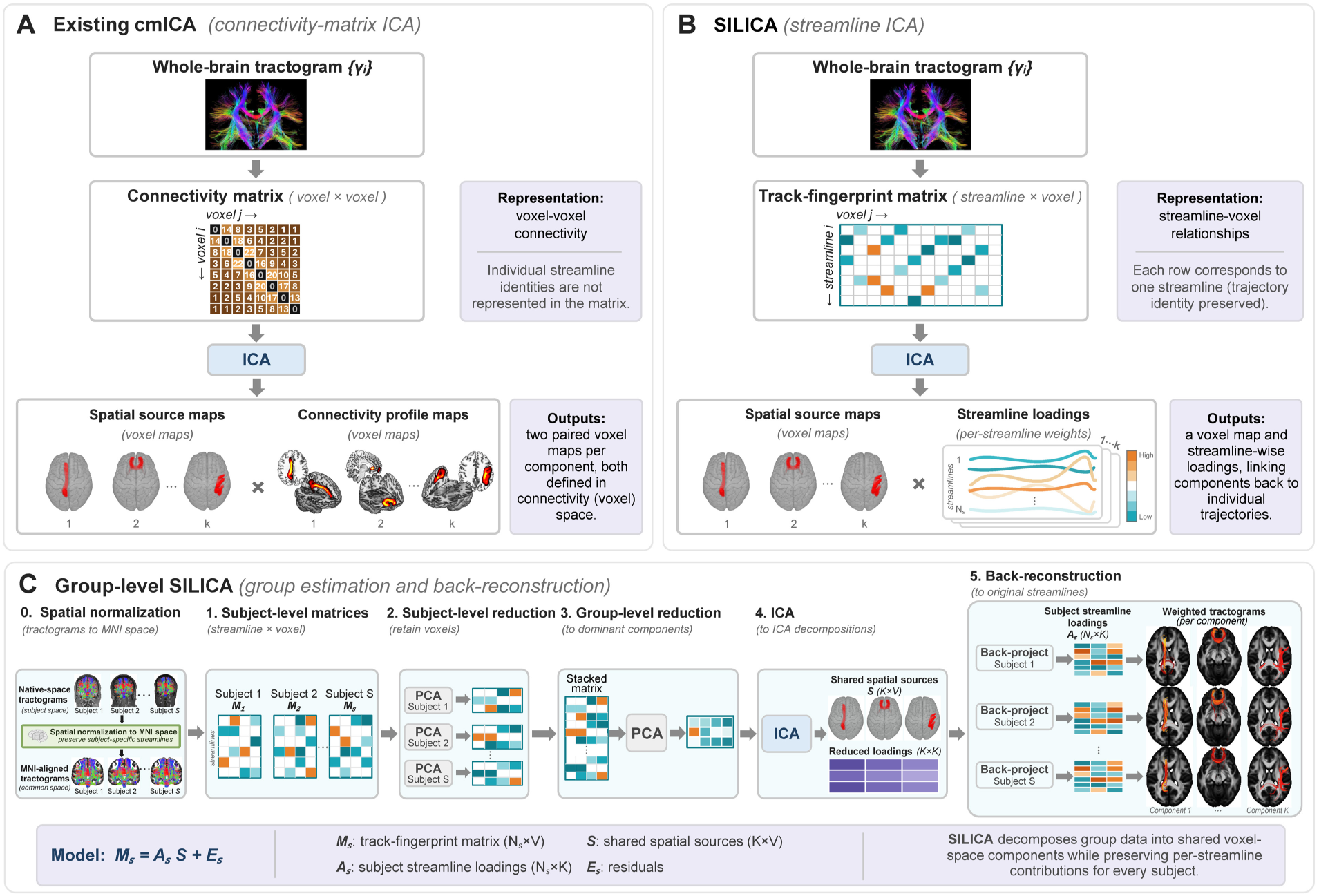
Overview of the SILICA framework. (A) Connectivity-matrix independent component analysis (cmICA). (B) Streamline independent component analysis (SILICA), which replaces connectivity-level observations with individual streamlines represented by their voxel-wise track fingerprints. (C) Group-level SILICA, including spatial normalization, subject- and group-level dimensionality reduction, ICA estimation, and back-reconstruction of subject-specific component loadings at the original streamline resolution.

The construction of the track-fingerprint matrices, within-voxel weighting, subject- and group-level dimensionality reduction, ICA estimation, component orientation, and back-reconstruction procedures are described in the following sections.

### 2.2 Study sample and diffusion MRI acquisition

We analyzed dMRI data from 30 healthy adult participants in the Center of Biomedical Research Excellence (COBRE) dataset acquired at the Mind Research Network. The sample had a mean age of 32.8 ± 5.6 years and included 15 women and 15 men. Imaging was performed on a 3 T Siemens TIM Trio scanner. Diffusion data were acquired with a single-shot spin-echo echo-lanar imaging sequence with a twice-refocused balanced echo, 30 non-collinear diffusion directions at b = 800 s/mm², five b = 0 images, repetition time = 9000 ms, echo time = 84 ms, field of view = 256 × 256 mm², acquisition matrix = 128 × 128, 72 slices, 2 × 2 × 2 mm³ isotropic voxels, one excitation, 3/4 partial Fourier encoding, and GRAPPA acceleration factor 2.

### 2.3 Preprocessing and whole-brain tractography

Data were prepreprocessed using FSL 6.0 as described in (Wu et al., 2015; Wu, Caprihan, Bustillo, Mayer, & Calhoun, 2018). Diffusion data were corrected for head motion and eddy-rrent distortions before tensor estimation. A diffusion tensor model was fitted to the preprocessed data (FSL 6.0), and whole-brain deterministic tractography was generated using CAMINO without tract-specific regions of interest (Mazumder, Kanyal, Wu, D. Calhoun, & Hye Ye, 2024; Mazumder, Wu, Calhoun, & Ye, 2025). Tracking used an anisotropy threshold of 0.2, a curvature threshold of 60 degrees, a Euler integration scheme, and a 0.5-mm step size. The resulting Camino streamline files were converted to TCK format using MRTRIX3 (www.mrtrix.org) to support spatial transformation, direct streamline-coordinate access for in-house MATLAB analysis, and visualization.

### 2.4 Spatial normalization of tractograms

To place tractograms within a common coordinate system, each subject’s fractional anisotropy (FA) image was nonlinearly registered to an MNI-space FA template (FMRIB58_FA_1mm.nii.gz) using ANTs (Avants et al., 2011). The corresponding point-transform chain was applied to every three-dimensional point along each streamline, thereby mapping the subject-specific tractogram into template space. Track fingerprints were constructed after transformation to template space so that each matrix column referred to the same voxel location across subjects.

This procedure established approximate voxelwise spatial correspondence while preserving the original streamline sampling and trajectory identities. No one-to-one or pointwise correspondence between streamlines from different subjects was required. Spatial normalization is summarized as the initial preprocessing step in the group-level SILICA workflow shown in Figure 1C.

### 2.5 Track-fingerprint representation and within-voxel weighting

Direct ICA of raw streamline coordinates is problematic because streamlines contain different numbers of sampled points. Resampling every streamline to a fixed number of coordinates produces equal-dimensional feature vectors but does not establish anatomical correspondence between coordinate indices. For example, a point with the same index along two resampled streamlines may correspond to different anatomical locations because the trajectories differ in length, curvature, orientation, and endpoint placement. A source defined over these coordinate indices may therefore be difficult to interpret anatomically, particularly in the context of a group analysis. SILICA instead represents each streamline by its occupancy within a common voxel coordinate system, in which every feature has a consistent spatial interpretation across streamlines and subjects.

Let γₛᵢ denote streamline *i* from subject *s*, and V the number of template-space voxels in the analysis mask. The track fingerprint mₛᵢ is a sparse vector in ℝ*^V^*, whose *v*th element records the within-voxel contribution of streamline *γγ_si_*:

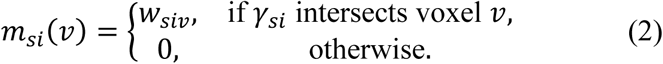

Stacking the streamline fingerprints yields the track-fingerprint matrix *M_s_* ∈ ℝ*^Ns^*^×*V*^. Each row represents one original streamline, and each column represents one template-space voxel. The matrix is sparse because each streamline intersects only a small fraction of the voxels in the analysis mask. Its nonzero pattern encodes the spatial course of the streamline, whereas the value *w_siv_* specifies how its contribution within voxel *v* is weighted.

Possible definitions of *w_siv_* include binary occupancy, the number of sampled segments within each voxel, and physical path length within each voxel. Binary occupancy records whether a streamline intersects a voxel but discards the extent of traversal. Segment counts depend on the density with which streamline points are sampled. Within-voxel path length provides a continuous geometric measure of trajectory occupancy that is less dependent on point-sampling density. In the present analysis, wₛᵢᵥ was defined as the physical path length of streamline γₛᵢ within voxel v.

### 2.6 Two-stage dimensionality reduction and streamline-level back-reconstruction

Following spatial normalization and construction of the subject-specific track-fingerprint matrices, SILICA used two stages of dimensionality reduction before group spatial ICA. First, each Mₛ was reduced via PCA independently from Nₛ streamline observations to rₛ = 100 subject-level components while retaining the common voxel dimension. The reduced subject matrices were concatenated along the observation dimension. Second, the concatenated matrix was reduced by PCA again to the selected group dimension K = 50 and decomposed using spatial ICA to get K group-level spatial components. This sequence accommodates unequal streamline counts across subjects and avoids constructing a single matrix that contains all original streamlines.

After ICA estimation, the group- and subject-level reduction transforms were propagated in reverse order to recover Aₛ at the original streamline resolution. Each reconstructed row remained indexed to the corresponding streamline in the subject’s transformed tractogram. Back-reconstruction therefore recovered component loadings without resampling, averaging, or otherwise modifying the streamline coordinates. The shared rows of S represent group voxel-space sources, whereas the columns of Aₛ represent each subject’s continuous streamline-level expression of those sources.

Component stability was evaluated with ICASSO by repeating the ICA estimation and clustering the resulting component estimates. Stability information was used to support descriptive component review rather than as external anatomical validation. The details can be repeated in Supplementary S1.

### 2.7 Component selection and anatomical interpretation

A 50-component model was estimated. Components were reviewed descriptively for spatial localization, predominance within white matter and visual correspondence with established commissural, projection, and association pathways. Components dominated by edge effects, ventricular patterns, spatially isolated termination patterns, or other structures inconsistent with white-matter anatomy were classified as obvious artifacts. Anatomical labels were assigned after estimation through visual comparison with established tractography atlases. These labels were used only for descriptive interpretation and were not incorporated as priors or constraints in the decomposition.

### 2.8 Component-specific representations

For subject *s*and component *k*, the corresponding column of *A_s_* was mapped back to the original streamline coordinates to generate a subject-specific component-weighted tractogram. Because the recovered loadings remain associated with the original streamlines throughout the back-reconstruction procedure, each trajectory can be visualized at its native three-dimensional coordinates without streamline resampling or geometric averaging. For visualization, streamline loadings were standardized separately for each subject and component, and only streamlines with *z* > 3were displayed. Streamlines were rendered with color proportional to the standardized loading. This threshold was applied solely for visualization and did not modify the continuous streamline loadings estimated by ICA. Selected components were visualized across 12 subjects to illustrate intersubject variation in trajectory geometry, spatial extent, streamline density, and anatomical dispersion.

Although the component-weighted tractograms preserve the original trajectory geometry, direct comparison across subjects remains limited by the absence of one-to-one streamline correspondence. To obtain a common image representation, the recovered streamline loadings were projected into template-space voxels using the generalized track-weighted imaging (TWI) framework of Calamante et al (Calamante, 2017), producing a track-weighted component map for each subject and component.

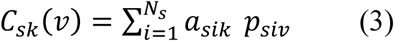

where *p_siv_* denotes the contribution of streamline *γγ_si_* to voxel *v* under the TWI projection. The recovered SILICA loading *a_sik_* therefore serves as the per-streamline weighting factor, and *C_sk_*(*v*) represents the accumulated contribution of component-weighted streamlines traversing voxel *v*. The projection was implemented efficiently as sparse matrix multiplication while preserving the generalized TWI formulation. Because these contributions are accumulated from continuously sampled streamline trajectories, the resulting maps inherit the super-resolution characteristics of the TWI framework. Direction-encoded (DEC) versions were additionally generated to visualize the local orientation of the contributing trajectories. Each subject and component therefore yields one track-weighted component map that provides a common image representation for visualization and future voxelwise statistical analysis.

### 2.9 Representative trajectory atlas

To provide a compact trajectory-space representation of the recovered components, a representative trajectory atlas was constructed from a group whole-brain tractogram. The MNI-registered diffusion tensor maps from all 30 subjects were combined to estimate a group diffusion tensor map, from which representative whole-brain tractography was performed. The resulting tractogram was encoded using the same track-fingerprint representation as the individual subject tractograms to obtain a group track-fingerprint matrix, *M*_grp_.

Rather than re-estimating the ICA model, the previously ICASSO-estimated group spatial components, *S*, were held fixed, and streamline-level component loadings for the representative tractogram were obtained by least-squares projection:

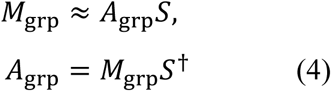

where *S*^†^ denotes the generalized inverse of *S*. The recovered streamline loadings were subsequently mapped to the corresponding trajectories in the representative tractogram, yielding one representative trajectory for each SILICA component. These trajectories provide a compact anatomical reference for visualization and interpretation of the group components while preserving the original SILICA component definitions.

## 3. Results

### 3.1 Spatial normalization and tractography variability

Visual inspection showed that transformation to MNI space aligned the overall location and orientation of the major white matter pathways across participants (Figure 2). Despite this common anatomical framework, the normalized tractograms retained visible inter-subject differences in streamline density, spatial dispersion, and local trajectory geometry. These preserved differences motivate the subsequent evaluation of whether a common spatial decomposition can recover consistent components while retaining subject-specific trajectory information.

**Figure 2.**
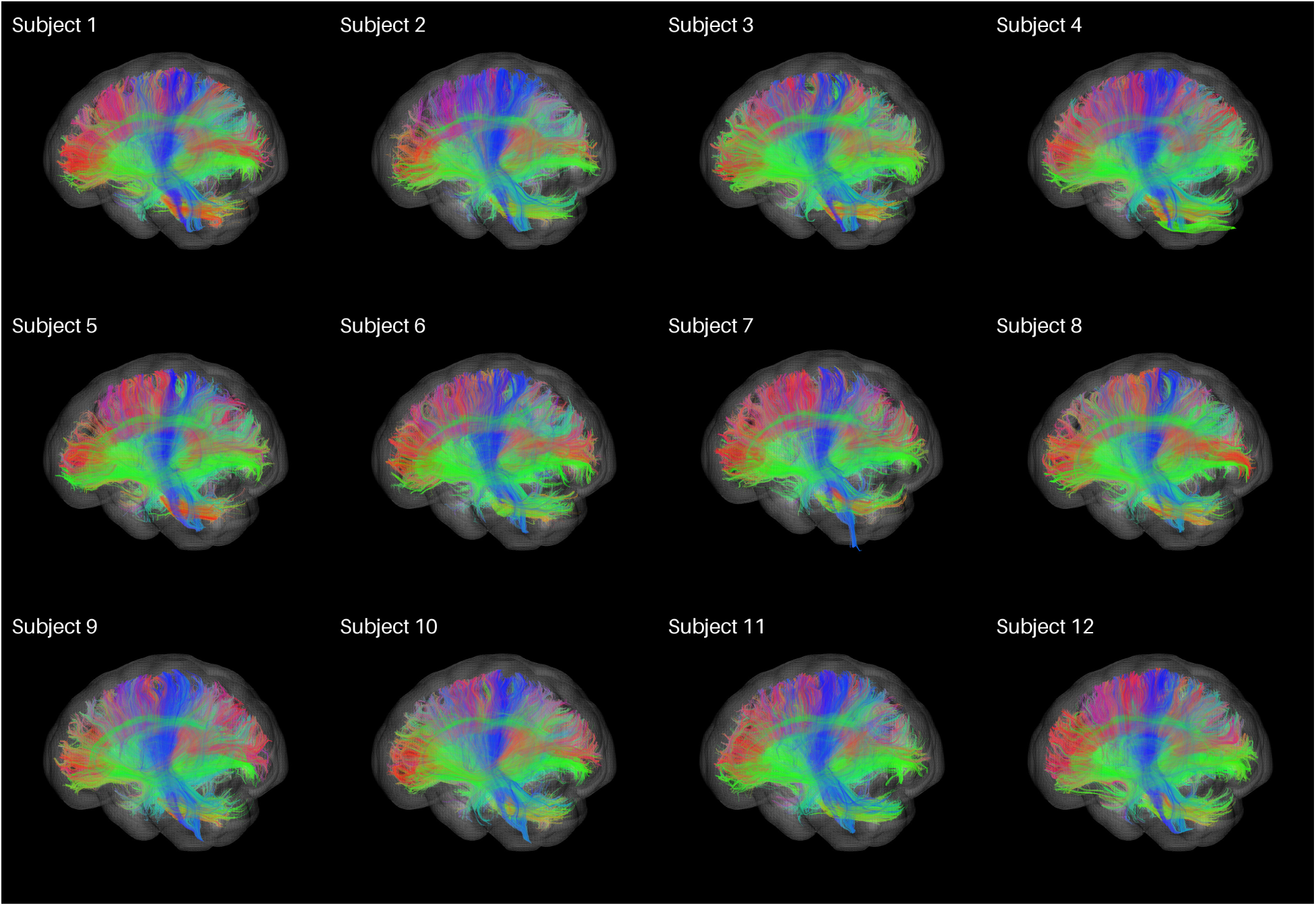
Representative MNI-normalized whole-brain tractograms from 12 subjects. Major white matter pathways occupy consistent anatomical locations following spatial normalization while preserving inter-subject differences in streamline density and local trajectory geometry. These normalized tractograms served as the input for subsequent SILICA decomposition.

### 3.2 Group-level spatial tract decomposition

Group SILICA estimated 50 independent components, of which 41 were retained following visual quality assessment (Figure 3). Nine components were excluded from further analysis as artifactual or anatomically indeterminate (Supplementary S2). Of these, five were dominated by cerebellar signal, while the other four exhibited overlapping, transitional, or spatially coherent patterns that could not be assigned a confident anatomical interpretation.

**Figure 3.**
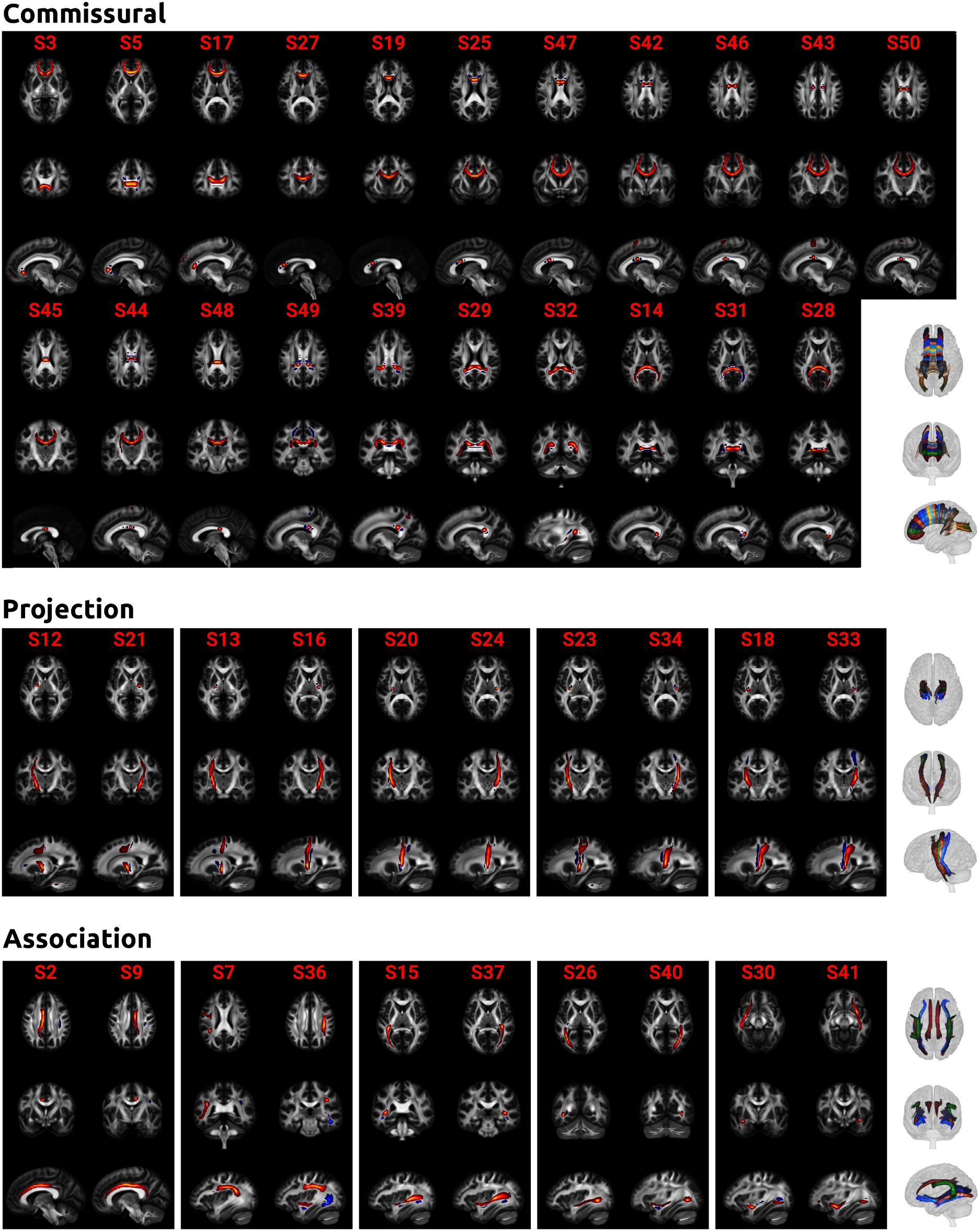
Group-level spatial source components from SILICA. Forty-one non-artifactual spatial components are organized into descriptive commissural (21 components), projection (10 components; five bilateral pairs), and association (10 components; five bilateral pairs) categories. Component labels were assigned after decomposition based on visual comparison with established white matter anatomy and are intended for descriptive interpretation only. Nine excluded components are shown in Supplementary S2.

The retained components exhibited spatial organizations broadly resembling major commissural, projection, and association white matter systems. Following our previous work (Wu & Calhoun, 2023; Wu et al., 2015), components were grouped according to conventional anatomical classifications (Mori, Wakana, Van Zijl, & Nagae-Poetscher, 2005; Wakana, Jiang, Nagae-Poetscher, van Zijl, & Mori, 2004) and guided by visual comparison with the Johns Hopkins University white-matter tractography atlas (Hua et al., 2008; Wakana et al., 2007). Commissural components showed predominantly midline and interhemispheric distributions, projection components followed superior-inferior orientations, and association components showed predominantly intrahemispheric distributions. These labels were assigned only after decomposition and were used solely for descriptive interpretation. No anatomical priors, tract labels, or tract-specific regions of interest were incorporated during component estimation, and anatomical correspondence was not quantitatively evaluated.

### 3.3 Subject-specific trajectory recovery

Back-reconstructed streamline loadings linked each group spatial component to the original tractogram of each participant, producing subject-specific weighted trajectory representations (Figure 4). Across the 12 representative subjects, the illustrated components followed broadly consistent anatomical courses while retaining visible intersubject differences in streamline density, spatial extent, curvature, and local trajectory geometry. The left and right cingulum and the forceps major and minor remained recognizable across participants, but their recovered streamline configurations were not identical, demonstrating that SILICA preserves subject-specific trajectory variation rather than imposing a single geometric template.

**Figure 4.**
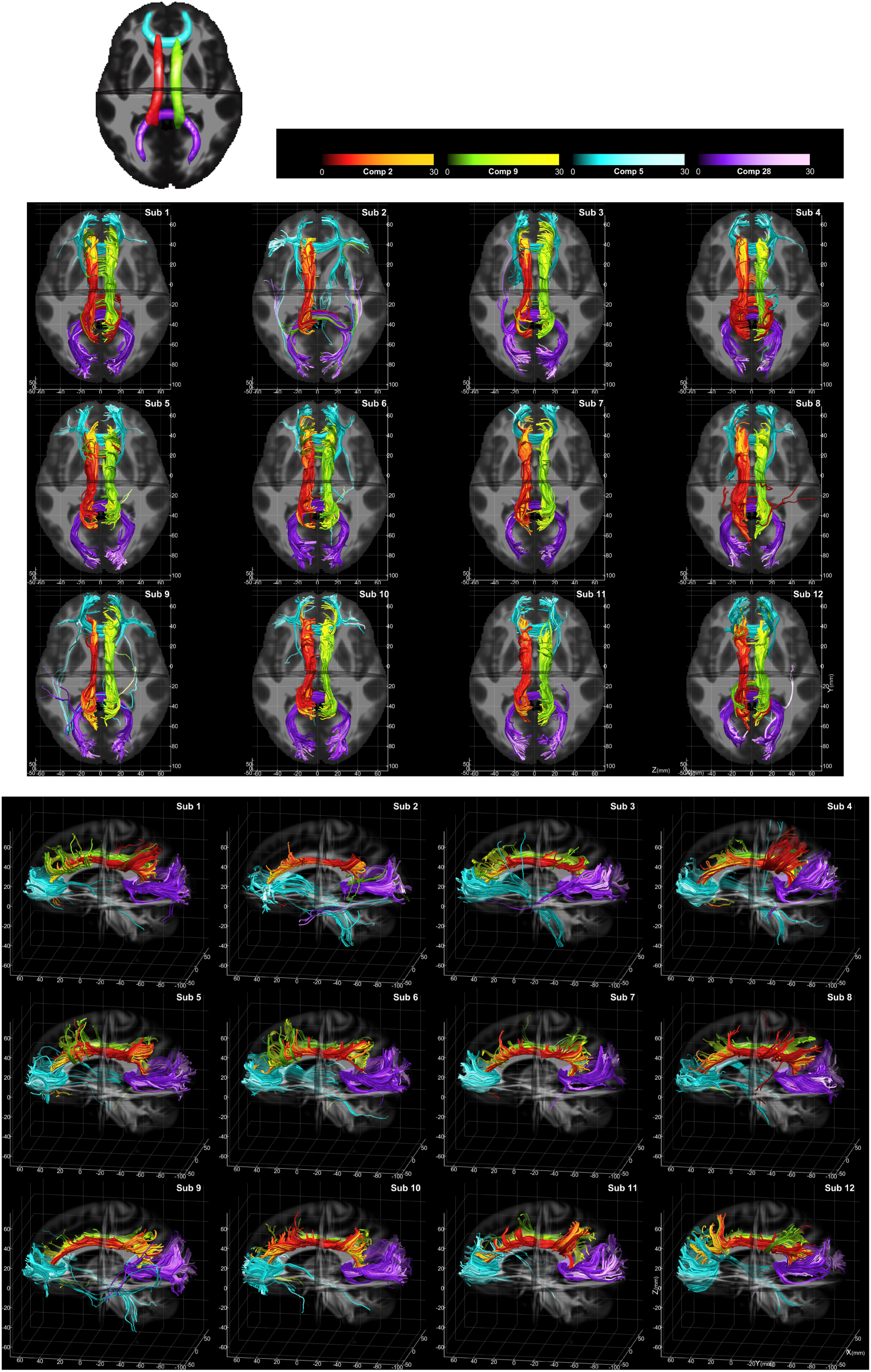
Subject-specific trajectory recovery and intersubject variability from SILICA components. Four representative components, left cingulum (red), right cingulum (green), forceps minor (cyan) and forceps major (purple), are shown across 12 subjects. The corresponding group spatial sources *S_k_* are displayed at the top, together with subject-specific weighted tractograms obtained by mapping *A_s_*(: ^,^ *k*) back to each participant’s original streamline coordinates. Axial superior and left sagittal views illustrate both the consistent anatomical course of each component and preserved intersubject variation in streamline density, extent, curvature, and local geometry. Streamline color intensity and opacity represent loading magnitude.

Mapping the oriented loadings to streamline color and opacity visualized the relative contribution of individual streamlines continuously, with higher-weight trajectories forming the dominant component core and lower-weight trajectories contributing more weakly. The resulting displays therefore represent graded component membership rather than binary tract segmentation. A single streamline could carry nonzero loadings for more than one component.

### 3.4 Track-weighted component maps

Projection of the recovered streamline loadings into template-space voxels using the generalized track-weighted imaging (TWI) framework produced subject-specific track-weighted component maps. Figure 5 shows the group-average maps across all 30 participants for the same four representative components shown in Figure 4, together with their directionally encoded color (DEC) representations. The group-average maps exhibited anatomically localized signal distributions corresponding to the recovered components, with higher voxel intensities reflecting the accumulated contribution of streamlines carrying larger component loadings, while the DEC maps depicted the local orientation of the contributing streamlines.

**Figure 5.**
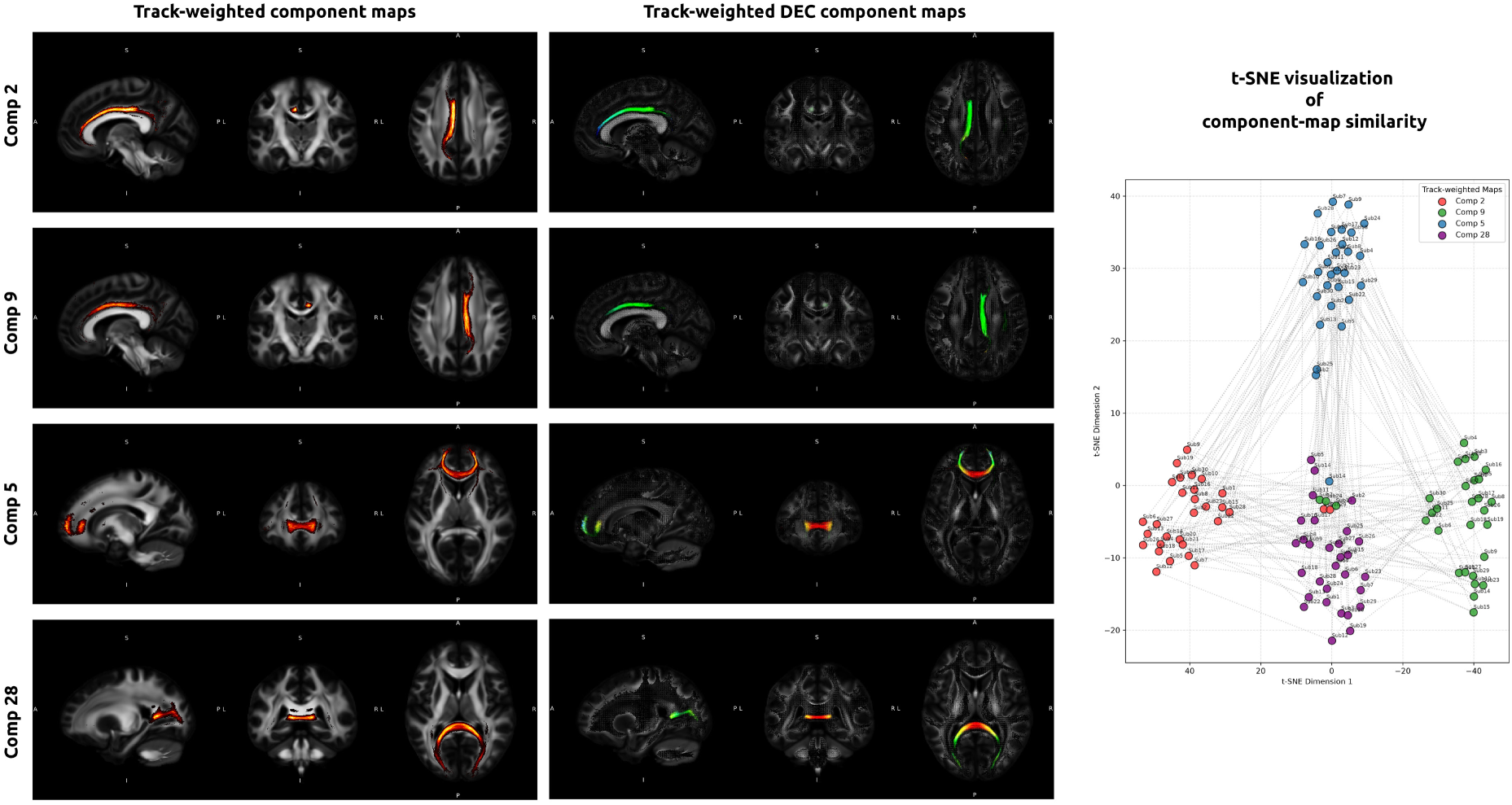
Group track-weighted component maps and t-SNE visualization. Group-average track- weighted component maps (left) and corresponding directionally encoded color (DEC) maps (center) for four representative SILICA components. Subject-specific maps were generated by projecting the recovered streamline loadings into template-space voxels using the generalized TWI framework and were then averaged across all 30 participants. In the DEC maps, red, green, and blue indicate left–right, anterior– posterior, and superior–inferior streamline orientations, respectively. Right: the exploratory t-SNE visualization of relative similarity among the subject-specific track-weighted component maps. Each color denotes one of the four illustrated components and follows the color convention used in Figure 4. The t-SNE embedding is provided for visualization only and does not constitute statistical clustering or validation of subject similarity.

This voxel-based representation provides a conventional image format for visualization and downstream voxel-wise analyses while preserving the continuous streamline weights recovered by SILICA. To illustrate how the subject-specific maps can support downstream quantitative exploration, t-distributed stochastic neighbor embedding (t-SNE) was used to visualize relative similarities among the track-weighted component maps in a two-dimensional space. This analysis was included as an exploratory visualization rather than a statistical test or validation of subject grouping. The maps were generated on a 0.5-mm isotropic output grid, producing the characteristic super-resolution appearance of TWI. As with conventional TWI, however, this apparent spatial detail reflects the accumulation of continuously sampled streamline trajectories on a fine voxel grid and should not be interpreted as increased biological resolution beyond the underlying diffusion MRI acquisition.

### 3.5 Representative trajectory atlas

To provide an intuitive group-level trajectory representation, the recovered SILICA spatial components were projected onto a representative whole-brain tractogram reconstructed from the group diffusion tensor template (Figure 6). Rather than averaging or selecting representative streamlines from individual subjects, the atlas was generated by decomposing an independently reconstructed whole-brain tractogram derived from the group diffusion tensor template. The resulting trajectories visualize the dominant pathway associated with each recovered component on a common anatomical substrate while preserving the continuous trajectory geometry of the representative tractogram and avoiding the need to establish one-to-one streamline correspondence across individuals. Representative trajectories were organized into commissural, projection, and association systems, providing an interpretable summary of the principal white matter organizations identified by SILICA. This trajectory-resolved atlas complements the voxel-space spatial components by providing a direct visualization of the corresponding three-dimensional streamline architecture.

**Figure 6.**
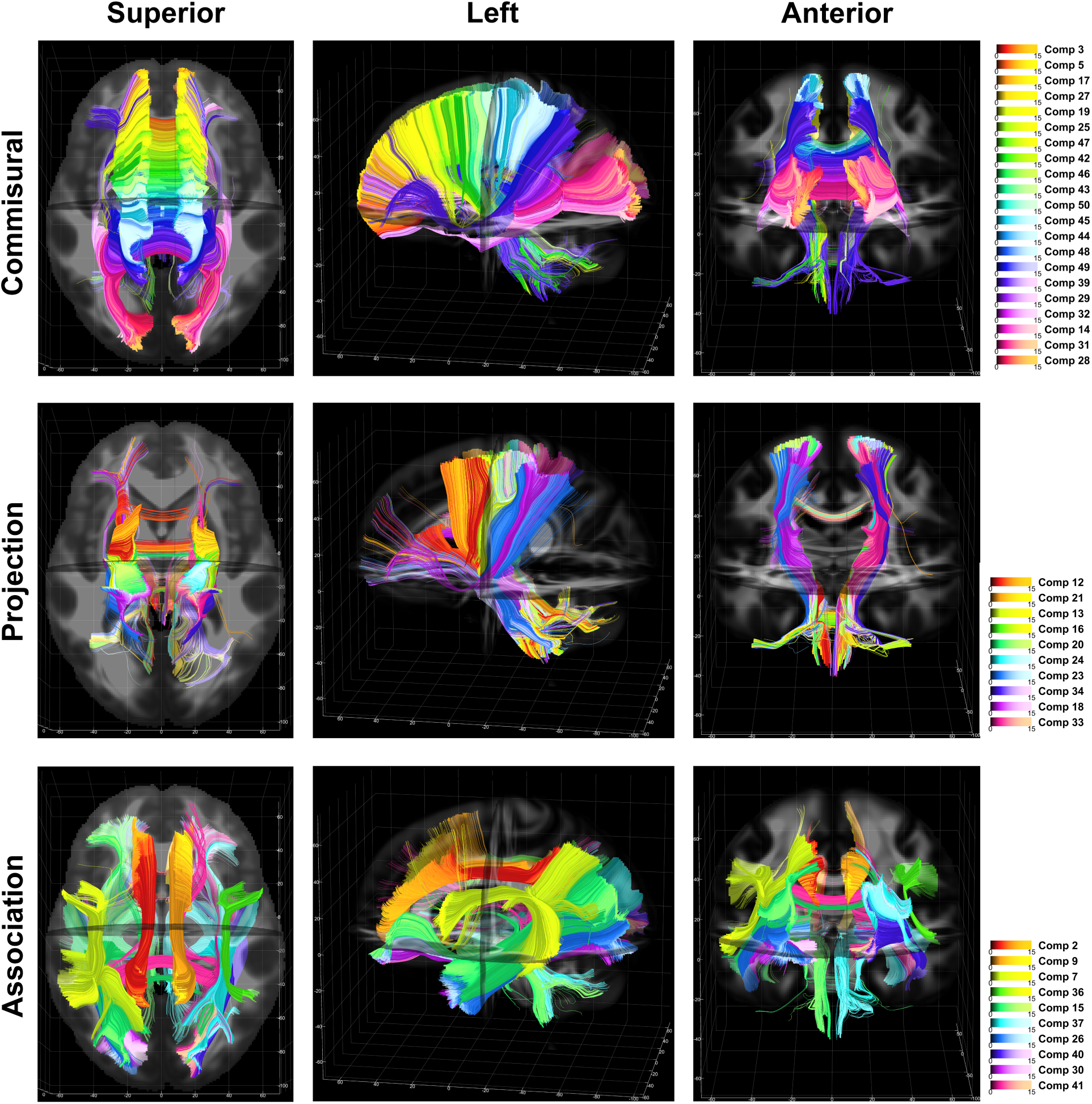
Representative trajectory atlas generated from SILICA. Representative trajectories were obtained by estimating streamline-level component loadings on a whole-brain tractogram reconstructed from the group diffusion tensor template while holding the recovered SILICA spatial components fixed. Components are organized into commissural, projection, and association systems according to the descriptive classification introduced in Figure 3. Each row shows superior (left), left sagittal (center), and anterior (right) views. Streamlines are colored according to their corresponding SILICA component. The atlas provides a compact group-level trajectory representation while preserving the continuous geometry of the representative tractogram.

## 4. Discussion

This proof-of-concept study introduces SILICA, a framework that links group-level voxel-space statistical decomposition with trajectory-resolved white matter representations through a sparse streamline-voxel fingerprint. The principal contribution is not simply the recovery of anatomically plausible white matter components, but the establishment of an explicit correspondence between statistical components estimated in voxel space and their associated streamline trajectories. Unlike standard ICA (on FA maps (Caprihan et al., 2011)) and our previous developed cmICA (on connectivity matrix), which produces only voxel-space component maps, or conventional streamline clustering, which operates exclusively in trajectory space, SILICA separates the statistical representation from the geometric representation while maintaining a direct mapping between the two. Consequently, each recovered component can be represented simultaneously as a spatial source, a subject-specific weighted tractogram, a track-weighted component map, and a representative group trajectory atlas.

Existing white matter analysis methods generally operate within either voxel space, streamline space, or connectome space. Streamline clustering preserves trajectory geometry but typically assigns discrete bundle membership, whereas connectome-based analyses summarize pathways as regional connections and largely discard their geometric organization. SILICA occupies an intermediate representation by preserving continuous streamline trajectories while assigning continuous component loadings within a common voxel-space basis. This representation naturally accommodates overlapping pathways and allows individual streamlines to contribute to multiple components without imposing hard tract boundaries. By maintaining an explicit correspondence between statistical and geometric representations, SILICA enables complementary analyses in voxel space, trajectory space, and track-weighted image space within a unified framework.

The sparse streamline-voxel fingerprint provides a common feature space without requiring one-to-one correspondence between streamlines across subjects. Following spatial normalization, each voxel represents a common anatomical location, whereas every streamline remains linked to its reconstructed trajectory. The accompanying two-stage dimensionality reduction further enables tractograms containing heterogeneous numbers of streamlines to be incorporated into a common group decomposition while maintaining computational tractability. The representative trajectory atlas presented here further illustrates that group-level trajectory representations can be generated without averaging streamlines or establishing streamline correspondence across subjects.

### 4.1 Tractography model and interpretation

The present data were acquired using a conventional single-shell diffusion protocol and reconstructed with a diffusion tensor tractography model. Because the tensor model estimates only a single principal orientation per voxel, it cannot resolve complex fiber configurations such as crossing, kissing, or fanning fibers. Consequently, the recovered components necessarily depend on both the diffusion acquisition and the tractography reconstruction model.

More fundamentally, SILICA operates on reconstructed tractograms rather than biological white matter itself. Streamline counts and streamline weights are influenced by acquisition quality, tractography parameters, seeding strategy, and reconstruction uncertainty, and therefore should not be interpreted as direct measures of axonal number or structural connectivity. Likewise, the recovered SILICA loadings quantify the contribution of reconstructed trajectories to each statistical component rather than biological connection strength. The derived track-weighted component maps therefore represent image-space visualizations of component-weighted tractography rather than quantitative measures of white matter connectivity. These limitations are common to tractography-based analyses and should be considered when interpreting the recovered components.

Although the present implementation uses tensor tractography, the SILICA framework itself is independent of the underlying tractography algorithm. Higher-angular-resolution acquisitions together with multi-fiber reconstruction methods, including constrained spherical deconvolution and other fiber-orientation-distribution approaches, can be incorporated without modifying the streamline-voxel representation or the decomposition framework.

### 4.2 Future directions and validation

The trajectory-resolved representation produced by SILICA enables several downstream analyses beyond those demonstrated in this proof-of-concept study. In voxel space, recovered streamline loadings can be projected into track-weighted component maps for conventional voxel-wise visualization and statistical analysis. In trajectory space, subject-specific loading matrices naturally support loading-weighted tractometry, streamline-based microstructural profiling, and quantitative measures of component expression across participants. More broadly, the common streamline-voxel representation provides opportunities for multimodal extensions, including joint decomposition with structural, functional, or molecular imaging, longitudinal analyses, and clinical group comparisons.

The present study is intended as an initial demonstration of feasibility rather than a comprehensive validation. External atlas comparisons, expert tract annotations, quantitative overlap metrics, test-retest reproducibility, systematic model-order analyses, and comparisons with streamline clustering and supervised tract segmentation remain important future directions. Additional evaluation across larger cohorts, higher-quality diffusion acquisitions, and alternative tractography algorithms will be necessary to establish robustness and generalizability. These limitations do not alter the central contribution of SILICA, a unified framework that estimates population-level statistical structure in a common voxel space while maintaining an explicit mapping to the subject-specific trajectories through which that structure is expressed.

## 5. Conclusion

SILICA introduces a trajectory-resolved ICA framework that links voxel-space statistical components to original subject-specific trajectories through a sparse streamline-by-voxel fingerprint. By maintaining an explicit correspondence between spatial component maps and their associated trajectories, the framework enables simultaneous visualization and analysis in voxel space, trajectory space, track-weighted image space, and representative group trajectory space without predefined tract labels or regions of interest. The proposed representation establishes a practical foundation for statistical comparison of component expression, loading-weighted tractometry, trajectory-resolved atlases, and multimodal decomposition.

## Acknowledgments

This work was supported by National Institutes of Health grants 1R01EB006841, 1R01EB005846, R01AG090597 and National Science Foundation grant 2112455.

## Ethics

All subjects provided written informed consent, and study procedures were approved by the Institutional Review Board at the University of New Mexico / Mind Research Network (MRN), under the COBRE program.

## Code and data availability

The COBRE dataset used in this study is available via the Collaborative Informatics and Neuroimaging Suite (COINS, https://coins.trendscenter.org/). Analysis code for the SILICA framework will be made available on GitHub upon publication.

## References

1. Avants, B. B., Tustison, N. J., Song, G., Cook, P. A., Klein, A., & Gee, J. C. (2011). A reproducible evaluation of ANTs similarity metric performance in brain image registration. Neuroimage, 54(3), 2033–2044.

2. Basser, P. J., Pajevic, S., Pierpaoli, C., Duda, J., & Aldroubi, A. (2000). In vivo fiber tractography using DT-MRI data. Magnetic resonance in Medicine, 44(4), 625–632.

3. Behrens, T. E., Woolrich, M. W., Jenkinson, M., Johansen-Berg, H., Nunes, R. G., Clare, S., . . . Smith, S. M. (2003). Characterization and propagation of uncertainty in diffusion-weighted MR imaging. Magn Reson Med, 50(5), 1077–1088.

4. Calamante, F. (2017). Track-weighted imaging methods: Extracting information from a streamlines tractogram. *Magnetic Resonance Materials in Physics*, Biology and Medicine, 30(4), 317–335.

5. Calamante, F., Tournier, J. D., Smith, R. E., & Connelly, A. (2012). A generalised framework for super-resolution track-weighted imaging. Neuroimage, 59(3), 2494–2503.

6. Caprihan, A., Abbott, C., Yamamoto, J., Pearlson, G., Perrone-Bizzozero, N., Sui, J., & Calhoun, V. D. (2011). Source-based morphometry analysis of group differences in fractional anisotropy in schizophrenia. Brain Connect, 1(2), 133–145.

7. Catani, M., & Thiebaut de Schotten, M. (2008). A diffusion tensor imaging tractography atlas for virtual in vivo dissections. Cortex, 44(8), 1105–1132.

8. Colby, J. B., Soderberg, L., Lebel, C., Dinov, I. D., Thompson, P. M., & Sowell, E. R. (2012). Along-tract statistics allow for enhanced tractography analysis. Neuroimage, 59(4), 3227–3242.

9. Garyfallidis, E., Côté, M. A., Rheault, F., Sidhu, J., Silva, D., Zhong, Q., . . . Descoteaux, M. (2018). Recognition of white matter bundles using local and global streamline-based registration and clustering. Neuroimage, 170, 283–295.

10. Hagmann, P., Cammoun, L., Gigandet, X., Meuli, R., Honey, C. J., Wedeen, V. J., & Sporns, O. (2008). Mapping the structural core of human cerebral cortex. PLoS Biol, 6(7), e159.

11. Hua, K., Zhang, J., Wakana, S., Jiang, H., Li, X., Reich, D. S., . . . Mori, S. (2008). Tract probability maps in stereotaxic spaces: analyses of white matter anatomy and tract-specific quantification. Neuroimage, 39(1), 336–347.

12. Jeurissen, B., Descoteaux, M., Mori, S., & Leemans, A. (2019). Diffusion MRI fiber tractography of the brain. NMR in Biomedicine, 32(4), e3785.

13. Johansen-Berg, H., & Behrens, T. E. J. (2009). Diffusion MRI: *From Quantitative Measurement to In Vivo Neuroanatomy*: Academic Press.

14. Mazumder, B., Kanyal, A., Wu, L., D. Calhoun, V., & Hye Ye, D. (2024). Physics-guided multi-view graph neural network for schizophrenia classification via structural-functional coupling. Paper presented at the International Workshop on PRedictive Intelligence In MEdicine.

15. Mazumder, B., Wu, L., Calhoun, V. D., & Ye, D. H. (2025). *Genetics encoded joint embedding of multimodal connectomes with explainable graph neural network for schizophrenia classification.* Paper presented at the 2025 IEEE 22nd International Symposium on Biomedical Imaging (ISBI).

16. Mori, S., Wakana, S., Van Zijl, P. C. M., & Nagae-Poetscher, L. M. (2005). MRI atlas of human white matter. Amsterdam: Elsevier.

17. O’Donnell, L. J., Golby, A. J., & Westin, C.-F. (2013). Fiber clustering versus the parcellation-based connectome. Neuroimage, 80, 283–289.

18. O’Donnell, L. J., & Westin, C. F. (2007). Automatic tractography segmentation using a high-dimensional white matter atlas. IEEE Trans Med Imaging, 26(11), 1562–1575.

19. Sporns, O., Tononi, G., & Kötter, R. (2005). The human connectome: A structural description of the human brain. PLoS Computational Biology, 1(4), e42.

20. Wakana, S., Caprihan, A., Panzenboeck, M. M., Fallon, J. H., Perry, M., Gollub, R. L., . . . Mori, S. (2007). Reproducibility of quantitative tractography methods applied to cerebral white matter. Neuroimage, 36(3), 630–644.

21. Wakana, S., Jiang, H., Nagae-Poetscher, L. M., van Zijl, P. C., & Mori, S. (2004). Fiber tract-based atlas of human white matter anatomy. Radiology, 230(1), 77–87.

22. Wasserthal, J., Neher, P., & Maier-Hein, K. H. (2018). TractSeg - Fast and accurate white matter tract segmentation. Neuroimage, 183, 239–253.

23. Wu, L., & Calhoun, V. (2023). Joint connectivity matrix independent component analysis: Auto- linking of structural and functional connectivities. Hum Brain Mapp, 44(4), 1533–1547.

24. Wu, L., Calhoun, V. D., Jung, R. E., & Caprihan, A. (2015). Connectivity-based whole brain dual parcellation by group ICA reveals tract structures and decreased connectivity in schizophrenia. Hum Brain Mapp, 36(11), 4681–4701.

25. Wu, L., Caprihan, A., Bustillo, J., Mayer, A., & Calhoun, V. (2018). An approach to directly link ICA and seed-based functional connectivity: Application to schizophrenia. Neuroimage, 179, 448–470.

26. Wu, L., Duda, M., Iraji, A., & Calhoun, V. D. (2025). Dynamic fusion of structural and functional connectivity via joint connectivity matrix ICA. bioRxiv, 2025.2005. 2015.653851.

27. Yeatman, J. D., Dougherty, R. F., Myall, N. J., Wandell, B. A., & Feldman, H. M. (2012). Tract profiles of white matter properties: Automating fiber-tract quantification. PLoS One, 7(11), e49790.

28. Yeh, F. C. (2020). Shape analysis of the human association pathways. Neuroimage, 223, 117329.

29. Zhang, F., Wu, Y., Norton, I., Rigolo, L., Rathi, Y., Makris, N., & O’Donnell, L. J. (2018). An anatomically curated fiber clustering white matter atlas for consistent white matter tract parcellation across the lifespan. Neuroimage, 179, 429–447.

