## Supplementary figures and images for "SILICA: Streamline Independent Component Analysis for Trajectory-Resolved White Matter Decomposition"

### Supplemental

**Supplemental**

S1. ICASSO
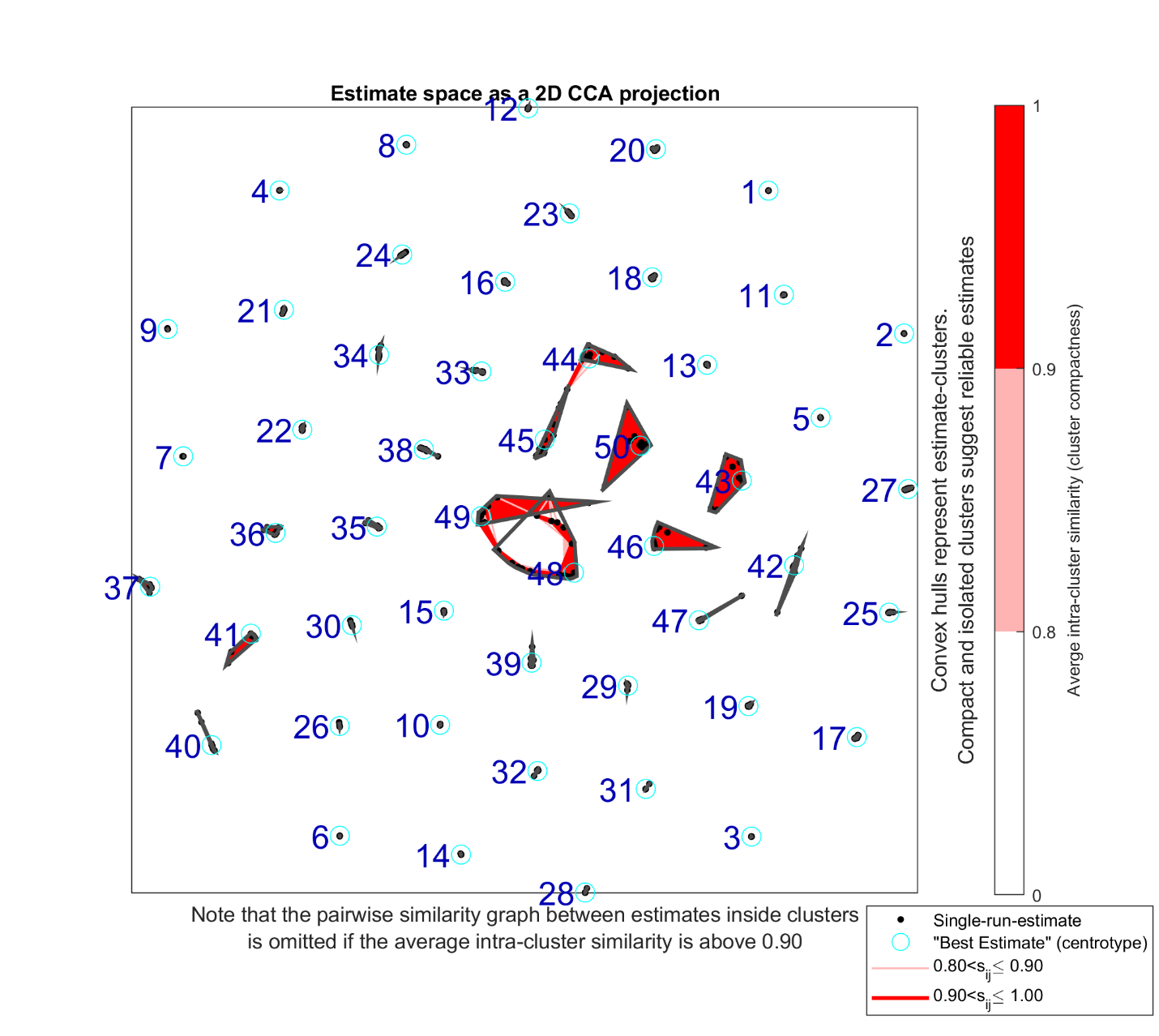

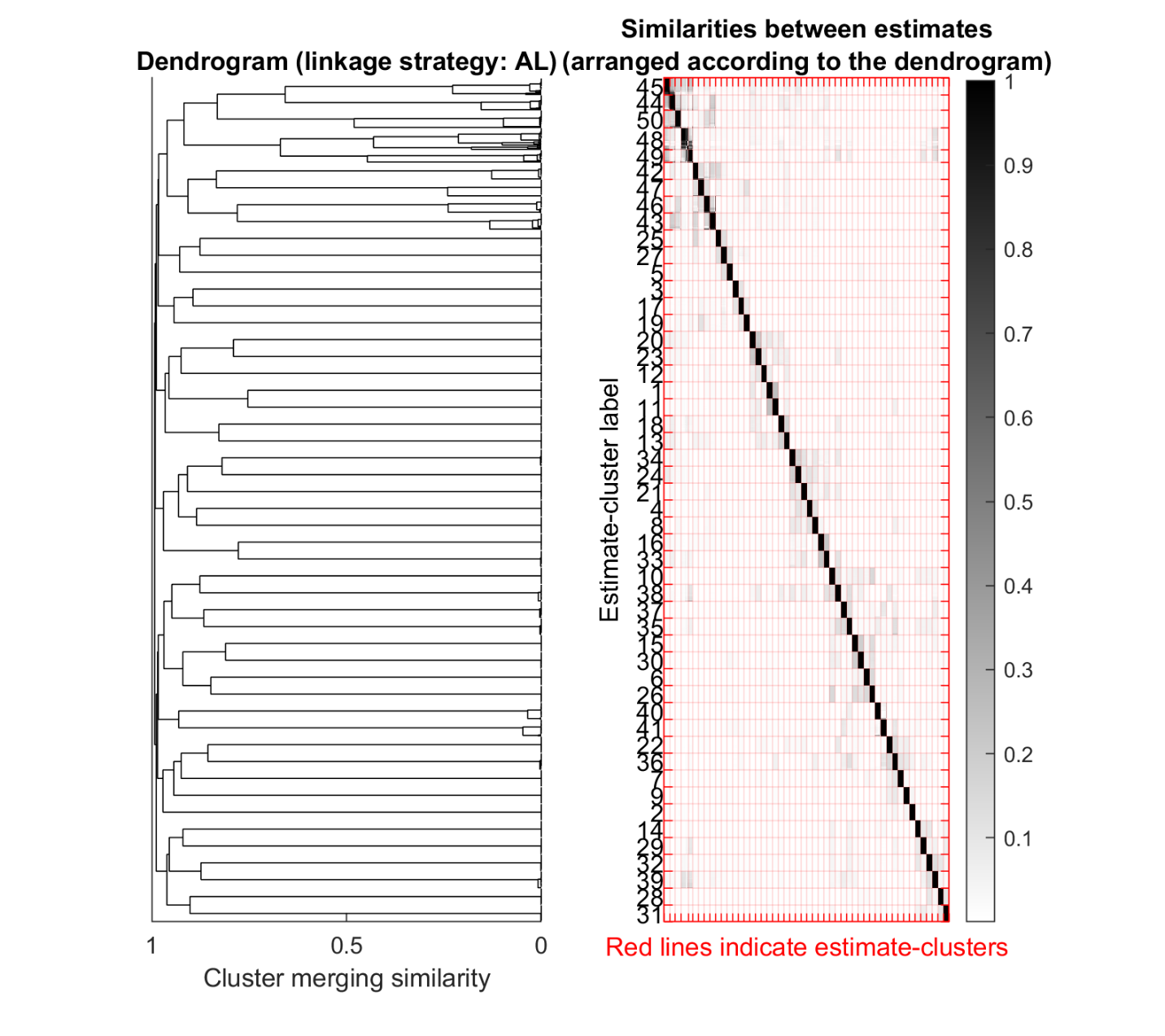

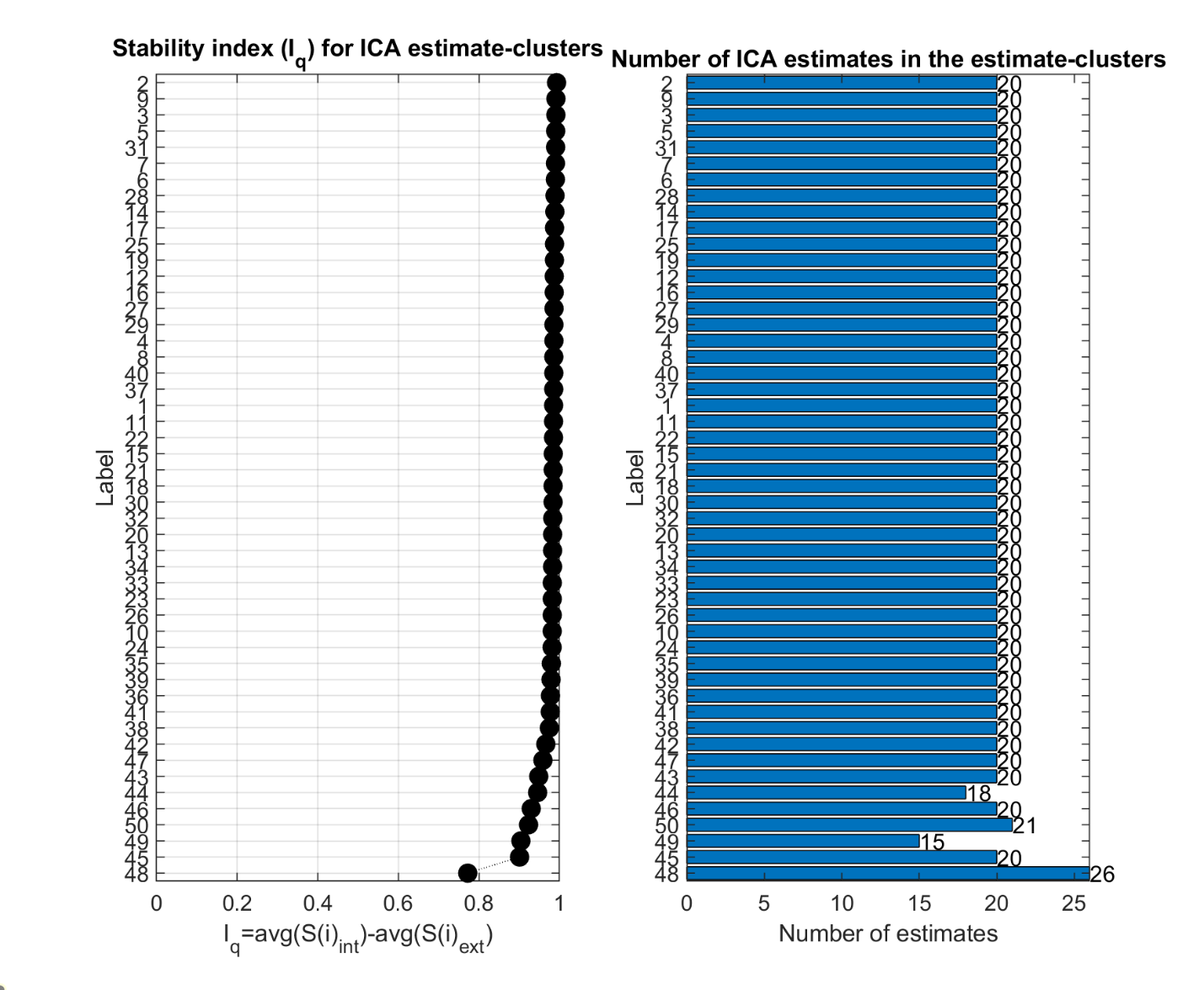

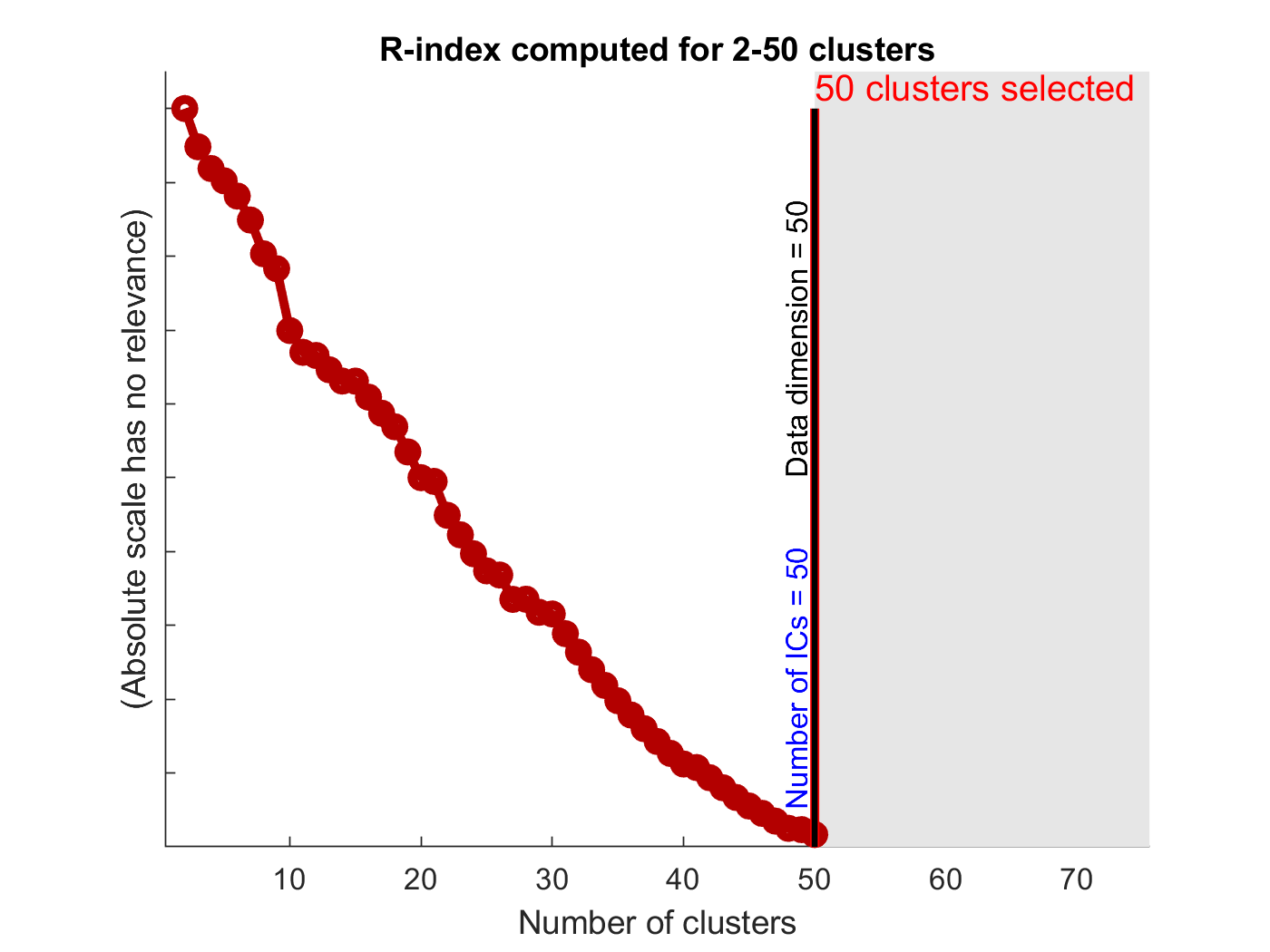

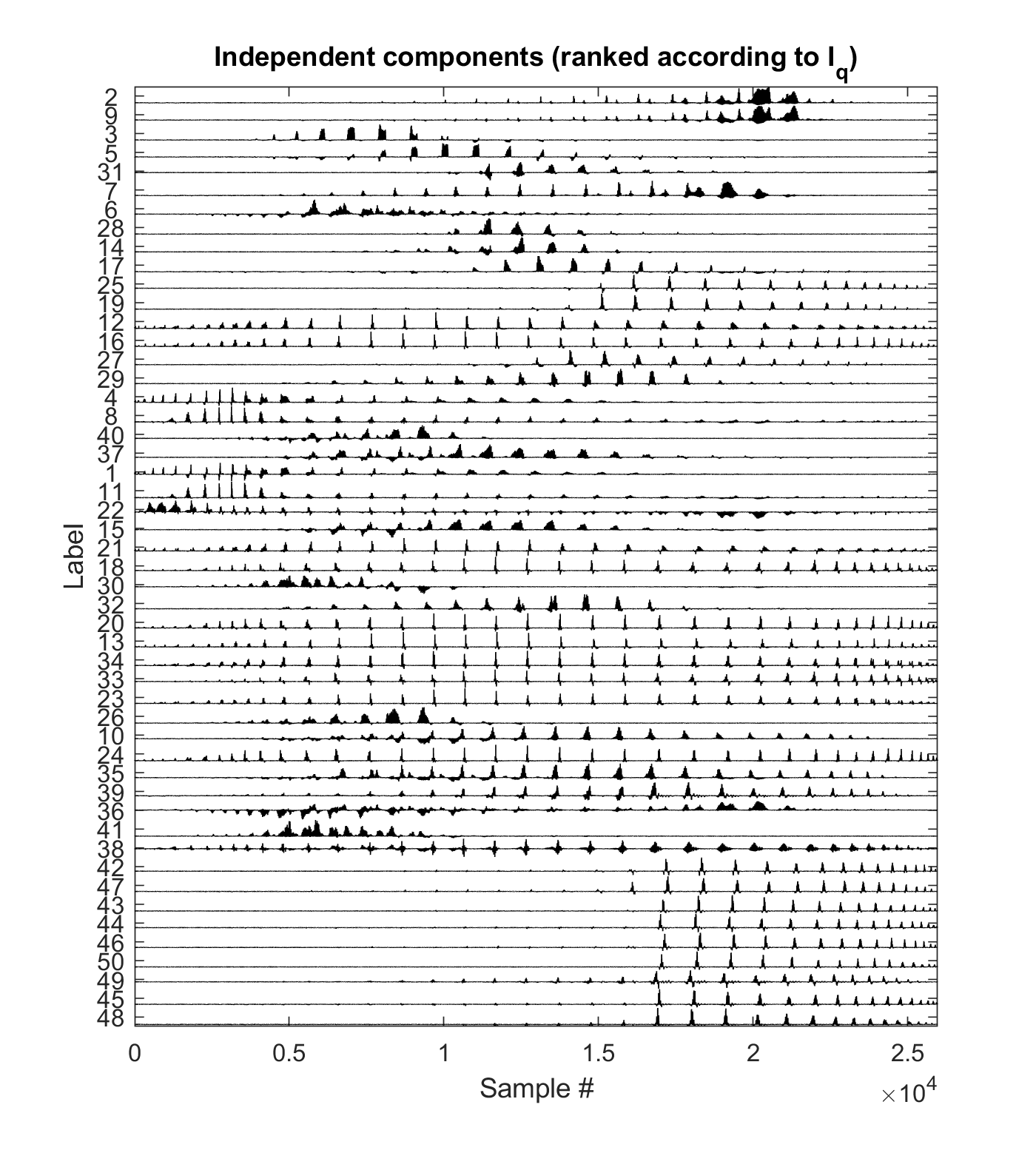

S2. Non-retained SILICA components

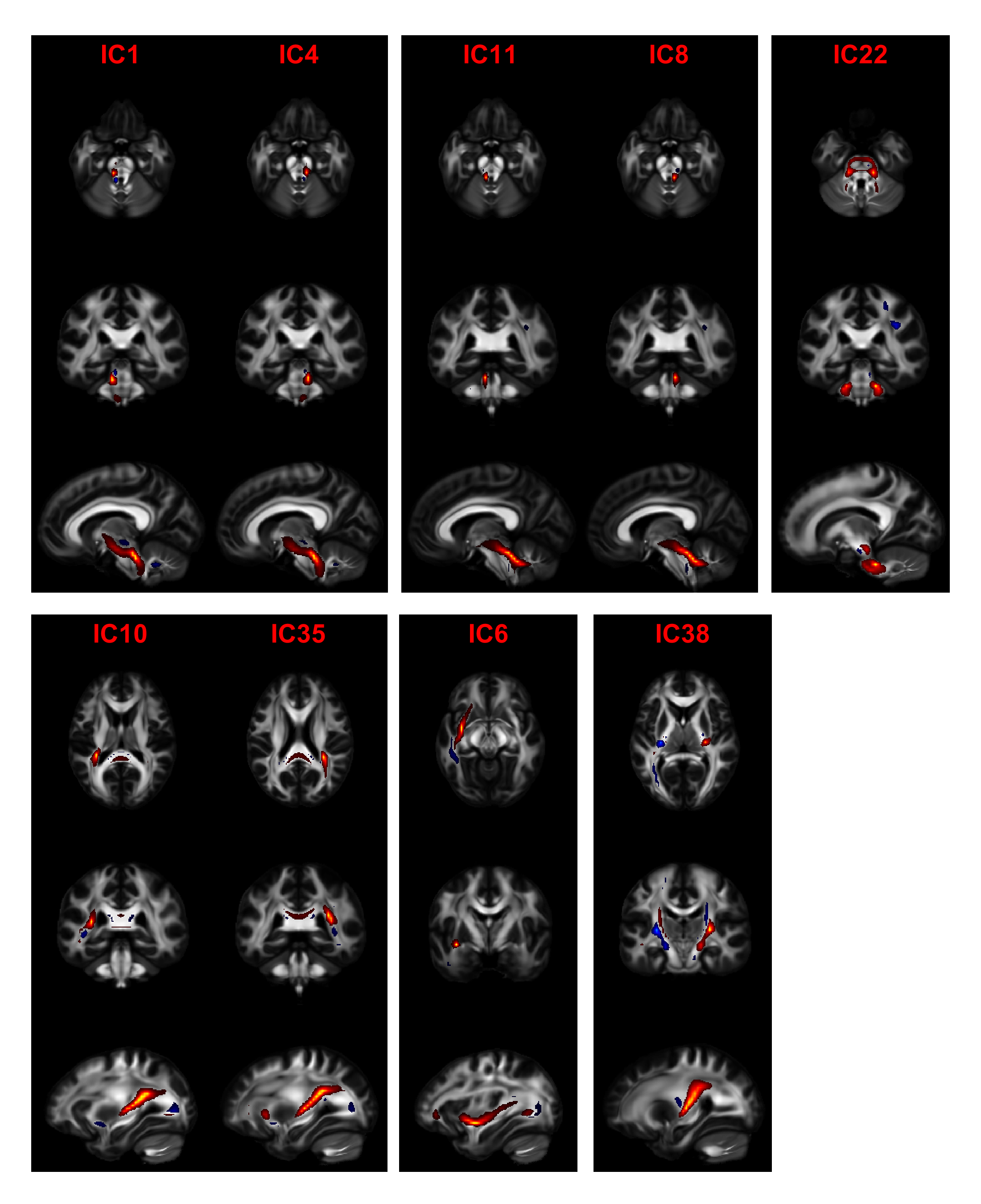
